# Vascular network-inspired diffusible scaffolds for engineering functional neural organoids

**DOI:** 10.1101/2024.08.31.610649

**Authors:** Hongwei Cai, Chunhui Tian, Lei Chen, Kyle McCracken, Jason Tchieu, Mingxia Gu, Ken Mackie, Feng Guo

## Abstract

Organoids, three-dimensional in vitro organ-like tissue cultures derived from stem cells, show promising potential for developmental biology, drug discovery, and regenerative medicine. However, the function and phenotype of current organoids, especially neural organoids, are still limited by insufficient diffusion of oxygen, nutrients, metabolites, signaling molecules, and drugs. Herein, we present Vascular network-Inspired Diffusible (VID) scaffolds to fully recapture the benefits of physiological diffusion physics for generating functional organoids and phenotyping their drug response. In a proof-of-concept application, the VID scaffolds, 3D-printed meshed tubular channel networks, support the successful generation of engineered human midbrain organoids almost without necrosis and hypoxia in commonly used well-plates. Compared to conventional organoids, these engineered organoids develop with more physiologically relevant features and functions including midbrain-specific identity, oxygen metabolism, neuronal maturation, and network activity. Moreover, these engineered organoids also better recapitulate pharmacological responses, such as neural activity changes to fentanyl exposure, compared to conventional organoids with significant diffusion limits. Combining these unique scaffolds and engineered organoids may provide insights for organoid development and therapeutic innovation.

## INTRODUCTION

Human brain organoids, 3D brain-like in vitro tissue cultures derived from human stem cells, can recapitulate key molecular, cellular, structural, and functional features of a developing human brain, holding remarkable potential for developmental biology, drug discovery, and regenerative medicine.^1-4^ So far, various neural organoids including cerebral organoids^5^, different region-specific organoids^6^, and assembloids^7^ have been generated for identifying novel mechanisms that underlie neurodevelopmental^8^, neurodegenerative^9^, and neuropsychiatric diseases^9^ as well as testing and screening compounds against these diseases^10^. Despite the tremendous promise, current human brain organoids still can’t fully recapitulate the physiology of a human brain in vivo, restricting their broader application for basic research and translational applications. One main limitation of current organoids with a diameter up to 3-4 mm is the lack of vascularization and circulation necessary to provide sufficient oxygen, nutrients, and growth factors and to eliminate waste during organoid differentiation.^3,11,12^ Consequentially, insufficient diffusion induces stressed, hypoxic, and necrotic cores in the interior of the spheroidal organoids and further affects the development and maturation of these organoids, possibly causing heterogeneity, reproducibility, and throughput issues.^13-15^

To address the diffusion limits of current brain organoids, pioneering attempts have been made by incorporating different perspectives.^16^ One perspective is to directly employ *in vivo* vascularization, perfusion, and circulation to support neural organoid development and function by transplanting human neural organoids or vascularized neural organoids into a rodent host, resulting in unprecedented survival of the organoid and even integration of organoid neural connectivity with the host brain. ^17,18^ An alternative solution is to leverage the organotypic slice culture at the air-liquid interface to enhance the diffusion of oxygen, nutrients, and growth factors for improving the survival and morphology of sliced cerebral organoids.^19,20^ Meanwhile, to overcome the diffusion limit to prevent cell death in the interior of organoids over long-term cultures, an advance has been made by culturing sliced neocortical organoids using a mechanical spinner, resulting in sustained neurogenesis and expanded cortical plate formation.^21^ Furthermore, engineering approaches such as organ chips^22-24^ and vessel chips^25-27^ have been developed to improve diffusion for engineering better organoids using sophisticated microfabricated devices^28-31^ and in vitro cultures of vascular networks^32-35^. However, challenges remain to develop simple and robust methods to support neural organoid development in a common lab setting.^1,11^

In a healthy human brain, each brain cell maintains excellent activity and function by receiving sufficient nutrients and oxygen supply through the physical diffusion of the vascular network.^36-38^ Here, we present Vascular network-Inspired Diffusible (VID) scaffolds (**Figure 1**) to fully recapture the diffusion efficiency of physiological vascular networks for generating functional neural organoids and phenotyping their drug response. The innovative scaffolds are designed by emulating the diffusion physics and biology of human brain tissues and are fabricated through the cost-effective, robust, and massive 3D printing of biocompatible plastics. These unique scaffolds can be simply incorporated into existing neural organoid protocols for generating engineered neural organoids with significantly reduced apoptosis, relieved stress, sustained neurogenesis, enhanced region-specific differentiation, and promoted function and drug response phenotype.

**Figure 1.**
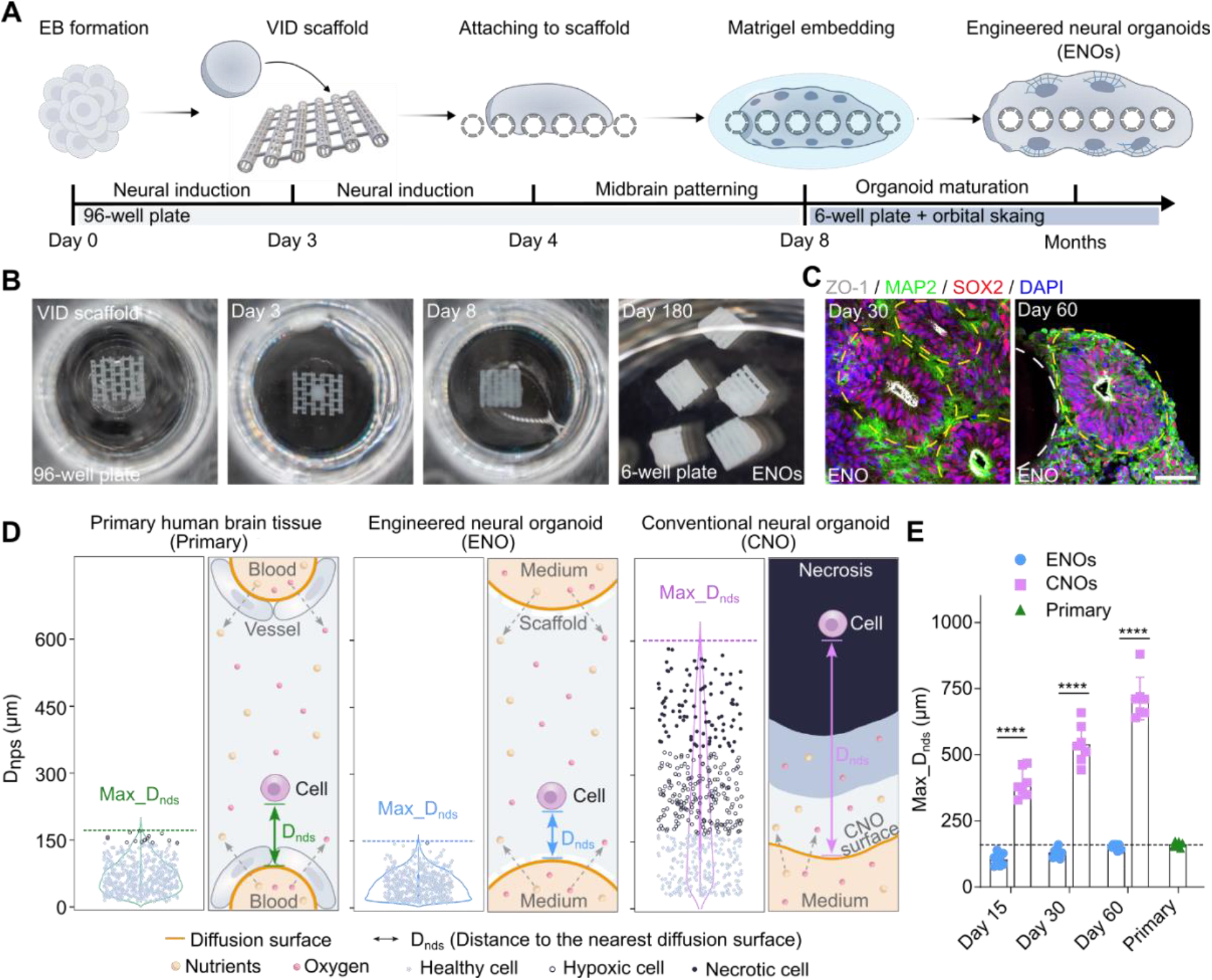
Generation of engineered neural organoids using VID scaffolds. **(A)** Schematic for generating engineered neural organoids (e.g., midbrain organoids) using Vascular network-Inspired Diffusible (VID) scaffolds within multiwell plates. **(B)** Representative bright-field images showing the engineered neural organoids at different time points in 96 well plates and 6 well plates. **(C)** Representative images of engineered neural organoids stained for ZO-1, SOX2, and MAP2 showing multiple ventricular zones (VZs). Yellow dashed lines indicate ventricular zones (VZs). Scale bars, 200 μm. **(D)** Distribution of healthy, hypoxic, necrotic cells along D_nds_ (distance to the nearest diffusible surface) in human primary brain tissues, engineered neural organoids (ENOs) at day 60, and conventional neural organoids (CNOs) at day 60. **(E)** Quantification of Maximum D_nds_ of the ENOs and the CNOs at day 15, day 30, and day 60, and Maximum D_nds_ quantified from human brain slides (In vivo) (mean± SEM, n = 6 from 3 independent experiments for ENOs and CNOs, n = 5 brain slides for primary tissues). White dashed circles indicate the VID scaffolds (panels **C**).

## DESIGN

Our objective is to develop engineered scaffolds that can recapitulate the diffusion physics of physiological vascular networks for transporting and releasing nutrients, oxygen, and therapeutics into, and removing wastes from, developing organoids. The successful development of the engineered scaffolds for generating functional organoids and improving their drug response phenotypes needs to consider several key criteria: (1) Engineered networks that recapture the diffusion function of vascular networks in vivo for maintaining the viability and function of all cells; (2) Biocompatibility and structural integrity for supporting the viability, development, and function of organoids over months; (3) Scalability, reproducibility, and cost-effectiveness that will broaden the use of organoids for basic research and translational applications; (4) User-friendly features such as compatibility with commonly-used well plates that can be easily adapted for various neural organoid protocols in common lab settings; and (5) Supporting various characterization methods for diverse applications of organoids.

To meet all the above criteria, we designed and fabricated simple, low-cost, robust, and compact VID scaffolds for high throughput generation and long-term culturing of flattened organoids with diffusible tube networks in well plates through 3D printing of biocompatible plastics. To determine scaffold design parameters, we investigated and compared how diffusion physics impacts the distribution of healthy, hypoxic, and necrotic cells in primary human brain tissues (primary tissues) and conventional neural organoids (**Figures 1D**). In the primary tissues, all brain cells are within 150 μm of a vessel wall, and they are healthy due to receiving sufficient nutrients and oxygen. In spheroidal midbrain organoids or conventional neural organoids (CNOs) at day 60, over 50% of the organoid cells are hypoxic or necrotic since they are about 150 μm to 600 μm away from the nearest organoid surface^39-41^. Thus, we define a parameter “D_nds_”, a cell’s distance to the nearest diffusible surface (e.g., vessel wall or organoid surface) for determining tissue diffusion. Moreover, to provide sufficient nutrients and oxygen (or avoid hypoxia and necrosis) for organoid culture, the maximum D_nds_ should be less than 150 μm^42^. Therefore, the VID scaffolds are designed to generate flattened neural organoids with diffusible tubal networks to keep the maximum D_nds_ of 150 μm for all organoid cells. Moreover, the scaffolds are expected to be compatible with the ubiquitous 96 well plates and to be easily handled via commonly used pipettes or tweezers in the lab. Thus, we propose a scaffold designed as a flattened matrix consisting of 6 parallel hollow meshed tubes.

## RESULTS

### Generation of engineered neural organoids using VID scaffolds

Based on the unique design features of the VID scaffold, we demonstrated their feasibility in generating flattened neural organoids with diffusible tube networks. We term these *engineered neural organoids (ENOs)*. As a proof-of-concept application, we incorporated the 3D-printed VID scaffolds with previously well-characterized midbrain organoid protocols (**Figure 1A**). This protocol involved the typical procedures including embryonic body (EB) formation, seeding EBs on scaffolds, midbrain patterning, Matrigel embedding, and organoid maturation. It is worth noting that the only added procedure is the seeding EBs onto biocompatible 3D printed VID scaffolds, while the other steps remain unchanged. The ENOs were generated as flattened midbrain organoids with diffusible tube networks (**Figure 1B**), and the successful development of these organoids was characterized by the formation of multiple ventricular zone (VZ) structures (**Figure 1C**). The distributions of healthy, hypoxic, and necrotic cells across D_nds_ were further analyzed in the ENOs at day 60, compared with those in the CNOs and primary human brain tissue (**Figure 1D**). The scaffolds with their flattened matrix design are expected to guide the flattened organoid formation to reduce the maximum D_nds_ during organoid development. The maximum D_nds_ in the developing ENOs at days 15, 30, and 60 were within 150 μm, similar to that in primary brain tissue, while the maximum D_nds_ in the developing CNOs at days 15, 30, and 60 increased from 394 μm to 720 μm (**Figure 1E**). These results showed that the maximum D_nds_ for all the organoid cells in the ENOs were maintained within 150 μm over two months of growth, and the cells were healthy almost without any hypoxic and necrotic conditions. This healthy growth is similar to that seen in the primary tissue but significantly different from that in the CNOs.

### Enhanced diffusion of medium, oxygen, and signaling molecules

The VID scaffolds are expected to employ the diffusion tube networks to directly supply medium-carrying nutrients, oxygen, and signaling molecules to the organoid cells. To validate these functions of our scaffolds during organoid development, we first characterized the perfusion of the medium by the scaffold (**Figure 2A**). The trajectories of the beads under the perfusion flow within the scaffolds indicated the transfer of fresh medium through the perfusable tube networks into the surrounding organoid tissues. Next, the supply of oxygen to the organoid during its development was also characterized numerically and experimentally. Hypoxia staining of ENOs at days 15, 30, and 60 found almost no hypoxia, while CNOs developed obvious hypoxic core and necrosis (**Figure 2B**), matching with numerical simulation results from the physics diffusion model. Quantification of hypoxia staining confirmed the same conclusion during the organoid development (**Figure 2C**). These results indicate sufficient oxygen transport for the ENOs during organoid development. Next, we demonstrated the perfusion of molecules with various molecular weights within the ENOs to determine if growth factors and other compounds would enter the ENO interstitium. In a perfusion assay using different molecules (Hoechst, 615Da; CF®488A WGA, 36kDa; CF@594 Con A, 105kDa) with orbital shaking, the results (**Figure 2D**) showed the enhanced and uniform fluorescent distribution of all three molecules across the ENOs, but in the CNOs, only a shallow fluorescent layer distributed on the surface was observed, indicating significantly enhanced perfusion and tissue penetration by the scaffold. The quantification of the dye penetration area in the ENOs and the CNOs confirmed this conclusion. Thus, we demonstrated that the perfusion provided by the scaffold can keep cells in a physiologically diffused-like condition without hypoxia and necrosis. With the well-characterized diffusion of our scaffolds, we next tested their biological effects.

**Figure 2.**
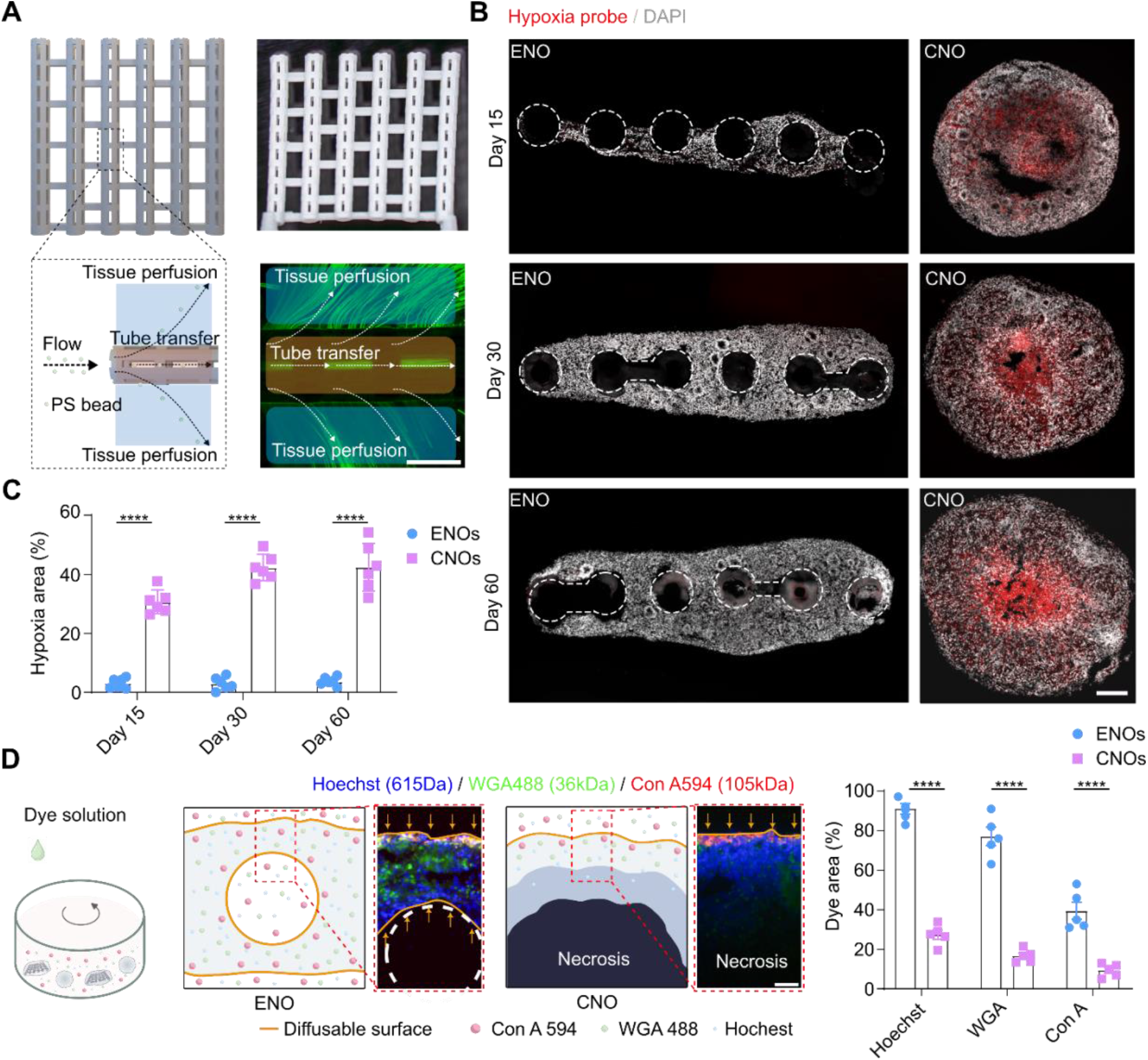
Enhanced diffusion of the medium, oxygen, and signaling molecules. **(A)** Top: schematics and bright-field image of the VID scaffold. Bottom: Schematics and overlayed trajectory of 2 μm beads showing the perfusion guided by VID scaffolds, including flow transfer in tubes and perfusion of the surrounding tissues. Scale bar, 200μm. **(B)** Pimonidazole-based hypoxia staining of the ENOs and the CNOs at days 15, 30, and 60. Scale bar, 200μm. **(C)** Quantification of hypoxia areas in the ENOs and the CNOs over time (mean± SEM, n = 6, from 3 independent experiments **(D)** Perfusion assay using different-sized dyes (Hoechst, 615Da; CF®488A WGA, 36kDa; CF@594 Con A, 105kDa) to model the tissue penetration of various molecules. Left: schematics and representative images showing dye distribution in the ENOs and the CNOs after 12 hours of perfusion. Scale bar, 50μm. Right: corresponding quantitative percentage of hypoxia areas in the ENOs and the CNOs (mean± SEM, n = 6 from 3 independent experiments). White dashed circles indicate the VID scaffolds (panels **B** and **D**).

### Relieved cell stress and sustained neurogenesis

Conventional neural organoids (CNOs) are not only subjected to significant hypoxia and necrosis but also suffer from broad cell stress and impaired metabolism compared to embryonic human brains, mainly due to insufficient diffusion of nutrients and oxygen.^13,43^ Consequentially, these abnormal cell conditions significantly affect neurogenesis and further limit the development and maturation of the organoids.^44^ By recapitulating physiological diffusion in the ENOs, we expect the limitations of CNOs will be mitigated with reduced cellular stress and substantially sustained neurogenesis using our scaffolds. To examine this hypothesis, we tested and compared developing ENOs and CNOs from day 15 to day 60, which is considered a key timeframe of progenitor expansion. We stained the ENOs and CNOs at days 15, 30, and 60 using a neural progenitor maker (SOX2), a mature neuron marker (MAP2), and an apoptosis marker (cleaved caspase-3, or Cas3) (**Figure 3A**). In addition, we stained for a proliferating cell marker (ki67) and an early-stage neuron marker (Tuj1) (**Figure 3B**). We found the ENOs and CNOs developed and matured over this time, including the emergence of mature neurons and the formation of VZs. By combining these observations with quantitative data (**Figure 3C**), we found that the CNOs started to present an obvious necrotic core (Caspase3+) at day 15, which continued to enlarge from day 15 to day 60, while the ENOs lacked a necrotic core and also showed less cell death in the non-necrotic proliferating region (**Figure S3A**) over the same time. Moreover, the ENOs maintained their neural progenitor ratio (measured by SOX2 staining) over time, but the CNOs decreased their neural progenitor ratio over time, despite starting with the same neural progenitor ratio at day 15. Furthermore, the ENOs developed a higher ratio of proliferating cells (ki67+) than the CNOs at day 60, although they had a similar level of proliferating cells (ki67+) at days 15 and 30.

**Figure 3.**
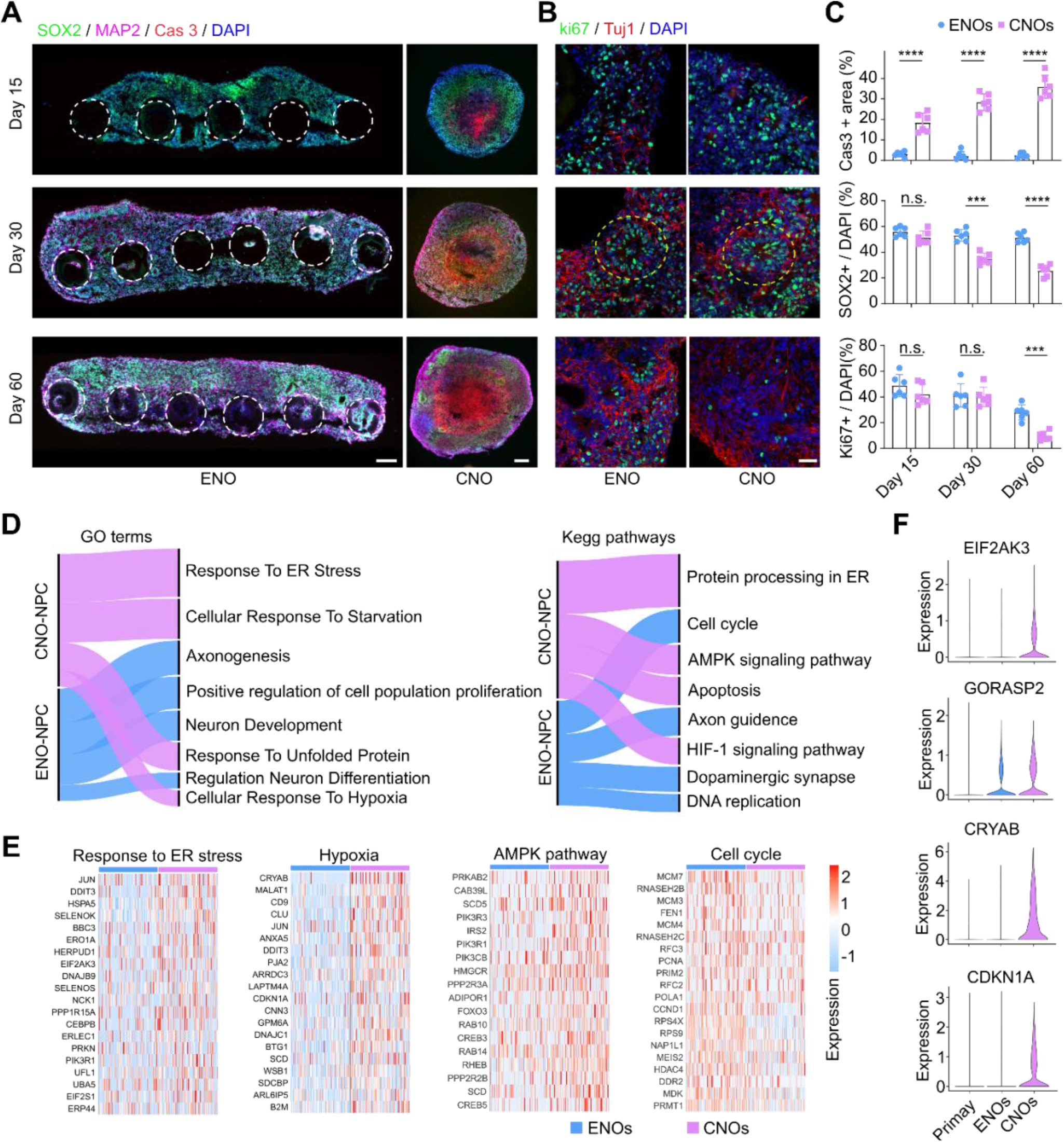
Reduced stress and sustained neurogenesis. **(A)** Representative immunostaining images of the engineered neural organoids (ENOs) and the conventional neural organoids (CNOs) at days 15, 30, and 60 for neural progenitor maker (SOX2), mature neuron marker (MAP2), and apoptosis marker (cleaved caspase-3, or Cas3). Scale bars, 200μm. **(B)** Immunostaining of the ENOs and the CNOs at days 15, 30, and 60 for proliferating cell marker (ki67) and early-stage neuron marker (Tuj1). Yellow dashed lines indicate ventricular zones (VZs). Scale bar, 200μm. **(C)** Percentage of neural progenitor cells (SOX2+), proliferating cells (ki67+) in the outer proliferating (non-necrotic) region at days 15, 30, and 60. Percentage of apoptotic area (Cas3+) in the whole organoid over time (mean± SEM, n = 6, from 3 independent experiments). **(D)** GO term and kegg pathway analysis of the most affected genes in the neural progenitor cells. Alluvial plots showing the selected gene ontology (GO) terms and kegg pathways comparing neural progenitor cells (NPCs) in the single cell RNA seq data of the ENOs and the CNOs at day 60. The thickness indicates the number of significantly different gene expressions involved in each term. **(E)** Heatmap of gene expression involved in GO term “Response to ER (endoplasmic reticulum) stress” and “Hypoxia”, kegg pathway term “AMPK pathway” and “Cell cycle” among neural progenitor cells (NPCs) in the ENOs and the CNOs from single-cell RNA seq data. **(F)** Expression of stress genes including EIF2AK3, CRYAB, GORASP2, and CDKN1A in the ENOs, the CNOs, and the primary human fetal midbrain tissues (primary) from single cell seq RNA seq data. White dashed circles indicate the VID scaffolds (panel **A**).

To comprehensively evaluate the status of neural progenitors, we performed scRNA-seq analysis of the ENOs and the CNOs on day 60. We analyzed the differently expressed genes of neural progenitor cells and performed downstream Gene Ontology (GO) and Kegg pathway analysis (**Figure 3D**). We found that upregulated genes in the progenitors from the CNOs were enriched in terms related to endoplasmic reticulum (ER) stress (response to ER stress, response to unfolded protein)^45,46^, hypoxia (cellular response to hypoxia, HIF-1 signaling pathway)^47^, impaired metabolism due to the lack of nutrients (cellular response to starvation, AMPK signaling pathway)^48^, and apoptosis (**Figure 3E**). Several genes involved in the cell cycle, DNA replication, and neuron development pathways such as UBE2C and STMN3 were significantly more expressed in progenitors from ENOs compared to their CNO counterparts. By further comparing our organoid data with a human embryonic midbrain dataset, we identified several genes involved in classical ER stress and impaired metabolism pathways that were highly expressed in the CNOs, such as EIF2AK3^49^, CRYAB^50^, GORASP2^13^, and CDKN1A, but these were expressed at significantly lower levels in the ENOs and human embryonic midbrains (**Figure 3F**). Together, compared to CNOs, our ENOs had reduced cell death, less cell stress, more highly supported cell metabolism, and sustained neurogenesis likely as a result of mimicking physiological diffusion physics.

### Promoted midbrain region-specific differentiation

Broad cell stress in conventional neural organoids is reported to impair cell-type specification^13,45^. With reduced cell death, relieved cell stress, and sustained neurogenesis, we expect the ENOs to have better midbrain differentiation, closer to authentic embryonic midbrain. To examine this, we first characterized and compared the proliferating midbrain progenitor cells in the developing midbrain proliferating zones in the ENOs and the CNOs at day 30 using LMX1A (dopaminergic progenitor marker) and ki67 (proliferating marker) (**Figure 4A**). A higher portion of LMX1A+ dopaminergic progenitors (46% vs. 23%) and ki67+/LMX1A+ proliferating dopaminergic progenitors (28% vs. 12%) were present in the VZs of the ENOs compared to those in the CNOs, as well as in overall organoids, indicating enhanced midbrain specific neurogenesis in the ENOs. Throughout organoid development, dopaminergic progenitors gradually transition to post-mitotic dopaminergic neurons expressing tyrosine hydroxylase (TH). The dopaminergic neurons (TH+) were observed surrounding the FOXA2+ (another dopaminergic progenitor marker) progenitors in the ENOs and the CNOs at day 60, but a higher ratio of FOXA2+ progenitors was found in the ENOs than in the CNOs (**Figure 4B**). Moreover, these TH dopaminergic neurons were predominantly colocalized with mature neurons expressing MAP2, and the ENOs consequently exhibited a higher proportion of dopaminergic neurons, as quantified by the ratio of dopaminergic neurons among mature neurons (37% vs. 25%) in the non-necrotic region (**Figure 4C**), as well as in overall organoids, compared to the CNOs. The gene expression (dopaminergic marker TH, dopaminergic progenitor marker NURR1, FOXA2, OTX2) detected by qPCR of the ENOs and the CNOs at day 60 further validated the enhanced midbrain-specific differentiation in the ENOs. Neuromelanin, previously reported as a sign of functional dopaminergic neurons in midbrain organoids^51-53^, could be observed as dark granular spots with more abundance in ENOs than those in CNOs.

**Figure 4.**
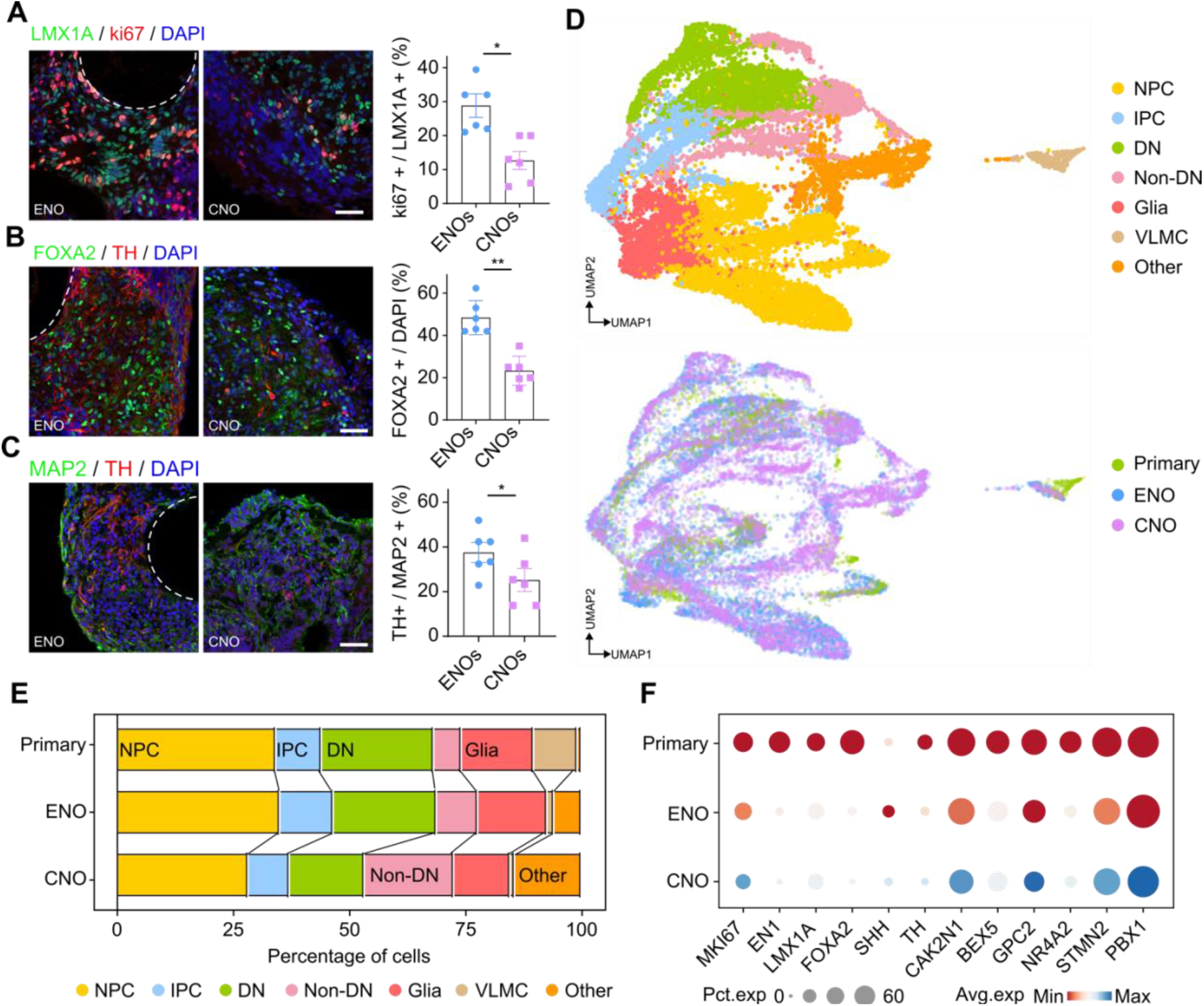
Enhanced midbrain region-specific differentiation. **(A)** Immunostaining of the engineered neural organoids (ENOs) and the conventional neural organoids (CNOs) at day 30 for dopaminergic progenitor marker (LMX1A) and proliferating cell marker (ki67). Scale bar, 200 μm. Corresponding quantification of proliferating dopaminergic progenitors (LMX1A+ & ki67+) (mean± SEM, n = 6 from 3 independent experiments). **(B)** Immunostaining of the ENOs and the CNOs at day 60 for dopaminergic progenitor marker (FOXA2) and dopaminergic neuron marker (TH). Scale bar, 200 μm. Corresponding quantification of dopaminergic progenitors (FOXA2+) (mean± SEM, n = 6 from 3 independent experiments). **(C)** Immunostaining of the ENOs and the CNOs at day 60 for mature neuron marker (MAP2) and dopaminergic neuron marker (TH). Scale bar, 200 μm. Percentage of dopaminergic neurons in ENOs and CNOs at day 60 (mean± SEM, n = 6 from 3 independent experiments). **(D)** Top: identification of various midbrain-like cell clusters in the integrated dataset. Uniform manifold approximation and projection (UMAP) plots of integrated single-cell RNA-seq dataset using canonical correlation analysis (CCA) method, 60-day ENO (9482 cells), 60-day CNO (8148 cells), and primary embryonic human midbrain (1977 cells). Bottom: UMAP plots showing the intersection of samples in the integrated dataset. Co-clustering of human primary embryonic midbrain and organoid single-cell RNA seq datasets. **(E)** Ratio of different cell clusters in the ENOs and the CNOs at day 60, and human primary embryonic midbrains. **(F)** Dot plot showing the expression of midbrain progenitor markers (EN1, LMX1A, FOXA2, and SHH) and dopaminergic neuron markers (TH, CAK2N1, BEX5, NR4A2, and PBX1) among three groups. White dashed circles indicate the VID scaffolds (panels **A, B**, and **C**).

To further unravel the differences of differentiation between the ENOs and the CNOs, we analyzed the cell types from the scRNA-seq data of the ENOs (9482 cells) and the CNOs (8148 cells) at day 60 and compared the results with a published primary human embryonic midbrain dataset (1977 cells)^54^ (**Figure 4D**). The organoid datasets were integrated with embryonic midbrain datasets using the canonical correlation analysis (CCA) method^55^, and cell clusters were identified using unsupervised clustering visualized on a Uniform Manifold Approximation and Projection (UMAP) plot. Based on previously reported embryonic midbrain marker genes, we assigned cell clusters to 7 distinct cell types, including progenitors, intermediate progenitors, dopaminergic neurons, non-dopaminergic neurons, glia, VLMC, and other unidentified cells. Pearson correlation analysis between the ENOs, the CNOs, and the primary human embryonic midbrain showed that cells in the developing human embryonic midbrain correlated to the corresponding cells in the ENOs, but not those in the CNOs. The cell composition analysis of these three conditions further demonstrated a higher similarity of the ENOs to the primary human embryonic midbrain (**Figure 4E**). Compared to the CNOs, reduced unidentified cells (5% vs. 14%) and non-dopaminergic neurons (8% vs. 19%), as well as increased dopaminergic neurons (22% vs. 15%) and progenitors (35% vs. 28%) were presented in the ENOs, which matched with our staining results. Moreover, we compared gene expression of midbrain-specific markers between the three datasets and found that the ENOs showed a higher expression level across all marker genes than CNOs (**Figure 4F**), indicating improved differentiation in the ENOs, closer to the primary conditions.

### Improved neural activity and network function

The multi-electrode array (MEA) systems are commonly used to measure and record neural activity of neural organoids for various applications^56,57^. However, there are several technological barriers to measuring functional neural spheroids, organoids, or 3D in vitro neural cultures: (1) Current MEA chips are not very flexible, and the detachment and low adhesion of 3D organoid cultures from the electrodes are common because the formation of physical contact between spheroidal cultures to the rigid and flat electrodes is not as easy as the adhesion of a 2D in vitro culture to the electrodes; (2) Many initially functional and active organoid neurons near the central MEA electrodes started to lose their neural activities and form low activity centers or neurotic cores due to the limited diffusion of this region after the organoid has been attached to the MEA chip for one- or two weeks. Given the flattened shape of engineered organoids guided by the VID scaffolds, we expect the ENOs to attach better to the MEA chips. Given the uniqueness of the scaffolds in recapitulating the physiological diffusion of nutrients and oxygen, we expect better neural activity and network function of the ENOs than the CNOs.

We characterized and compared the neural function of the ENOs and the CNOs at day 90 using MEA extracellular recordings. The organoids were attached per well in a 6-well MEA plate, containing 64 electrodes per well, 4 weeks before electrical activity measurements. During the attachment of the ENO onto the MEA chip, the unique flattened shape of ENOs perfectly conformed to the flat MEA electrodes, and the VID scaffolds continued to diffuse organoids with nutrients and oxygen to support energy-consuming neural network functions, however, the diffusion was limited in the CNOs, especially in the region near to the MEA electrodes (**Figure 5A**). Both the ENOs and CNOs showed spontaneous activity recordings, periodically occurring network bursts, in which wide-spreading bursts appeared across electrodes simultaneously indicating the maturation of the organoid functional networks (**Figure 5B**)^58^. The ENOs exhibited evenly distributed robust neural activity across the MEA sensing area, whereas CNOs displayed a distinct necrotic core with no detectable electrical activity (**Figure 5C**). The functional connectivity of the ENOs and the CNOs was thoroughly analyzed from the correlated spontaneous network activity (**Figure 5D**). A well-defined neuronal network was found in the ENOs, in which densely connected local networks could be found with both short- and long-range connections between electrodes. In contrast, a considerably sparser neuronal network was present in the CNOs, especially with reduced long-range connections, likely caused by the necrotic core. Through quantitative analysis of spontaneous activity, we found the ENOs showed significantly better neural activities than the CNOs through the quantified active electrodes (42.5 vs. 24.3), burst frequency (0.52Hz vs 0.21Hz), and mean firing rate (3.12Hz vs 1.05Hz) (**Figure 5E**). Moreover, the ENOs have significantly enhanced functional connectivity as quantified by classical network index (e.g., average clustering, local efficiency, global efficiency, and network density), compared to the CNOs. Additionally, analysis of the detected spikes revealed that the ENOs exhibited significantly higher amplitude spikes (**Figures 5F, G**), suggesting a more advanced maturation state of neurons^59,60^. Thus, we have demonstrated that our scaffolds improved neural activity and network function in the ENOs compared to the CNOs.

**Figure 5.**
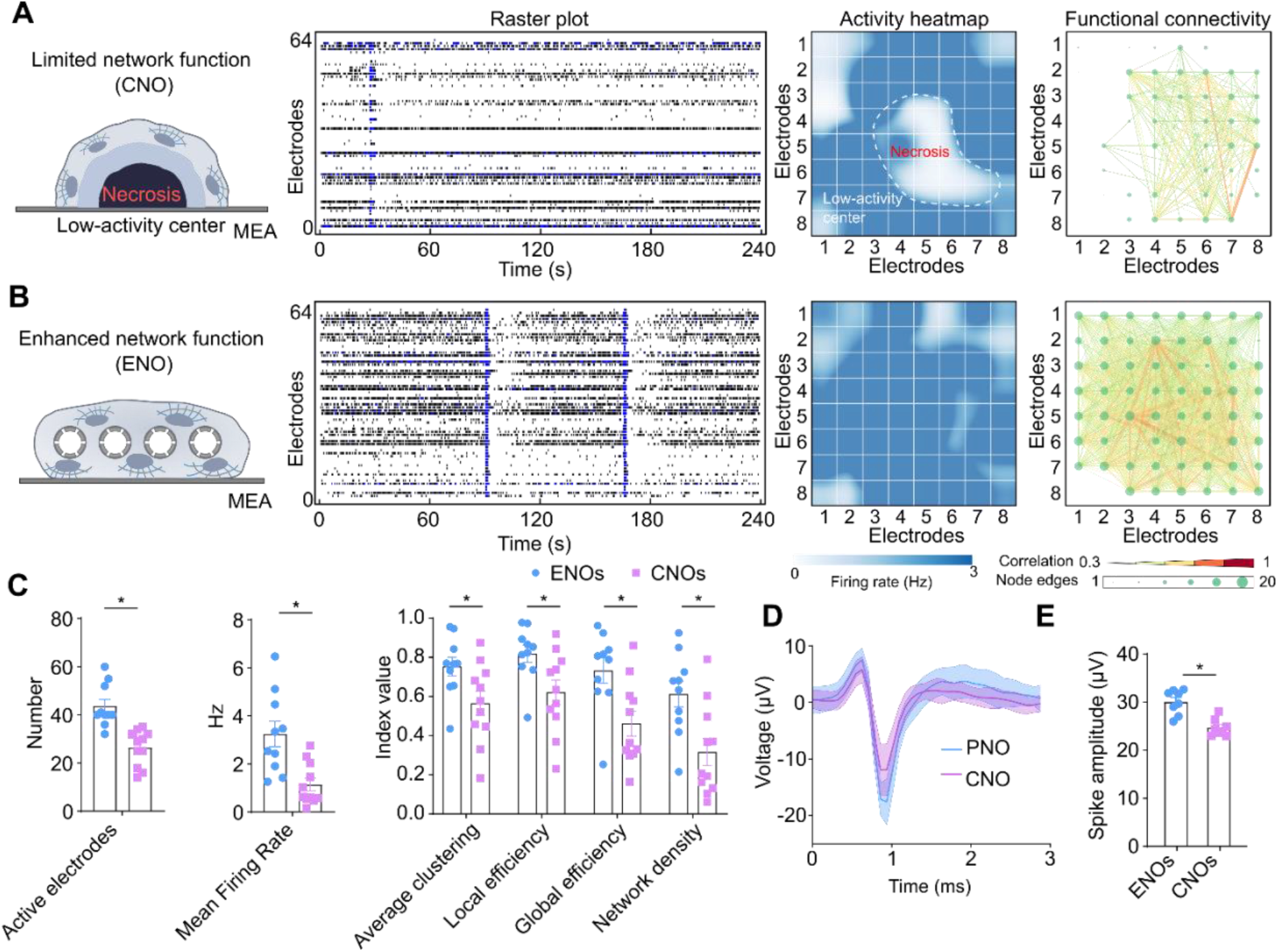
Improved neural activity and functional networks. **(A)** Schematics of measuring neural activity of conventional neural organoid (CNO) on MEA electrodes. A necrotic core with low neural activity was formed in CNOs as the diffusion from the MEA side was limited. Representative raster plots and smoothed activity heatmaps showing the network spiking activities in the CNO at day 90, a low-activity center could be observed in the CNO. Corresponding calculated functional connectivity maps. **(B)** Schematics of measuring neural activity of engineered neural organoid (ENO) on MEA electrodes. Enhanced diffusion by the VID scaffold could support the viability and function of the ENOs on MEA. Representative raster plots and smoothed activity heatmaps show the network spiking activities in the ENO at day 90, with good activity all over the organoid. Corresponding calculated functional connectivity maps. In the functional connectivity map, the color and thickness of the lines (edges) indicated the correlation level between electrodes, and the size of the nodes showed the number of lines (edges) connected to the corresponding electrode. **(C)** Quantification of active electrodes, mean firing rate, and network indexes (efficiency, density, and clustering of the functional network) of the ENOs and the CNOs at day 90 (mean± SEM, n = 11 from 3 independent experiments). **(D)** Representative spikes detected from one channel (with the highest spiking amplitude) of the ENO and the CNO. Solid traces represent the averaged spikes (30 spikes overlayed), and the ranges represent the SEM. **(E)** Quantification of spiking amplitudes from the ENOs and the CNOs, each point represents the averaged spiking amplitude detected from the electrode with the largest spiking amplitude in an organoid (mean± SEM, n = 8, from 3 independent experiments).

### Enhanced pharmacological responses

Measuring neural activity response of organoids using MEA holds promising potential for high throughput screening of compounds to identify novel treatments for various neurological disorders. However, current conventional organoids face two significant challenges (**Figure 6B schematics**): (1) the neurons near the MEA electrodes lose their electrical activity due to the impact of hypoxic and neurotic conditions, and (2) the drugs can’t easily reach the healthy neurons adjacent to the MEA electrodes or only access a small fraction of them. Given that our scaffolds can enhance organoid functionality and conform with the flat MEA electrodes to address the first issue, we further validated the ability of the ENOs to address the second challenge in the functional phenotyping of organoids for drug response by leveraging the excellent diffusion of the scaffolds. We used fentanyl, an effective analgesic widely used to relieve pain^61,62^, as a proof-of-concept demonstration for functional phenotyping of acute drug response. The spontaneous activity of the same ENOs was recorded before and 20 minutes after treatment with a clinically relevant concentration of fentanyl (50nM). The results showed obvious pharmacological effects including a reduction in the number of spike events and bursts and a subsequent loss of synchronous network activity (**Figure 6A**). In contrast, the CNO’s electrical activity showed a non-significant response to fentanyl (**Figure 6B**). Moreover, detailed quantifications including mean firing rate, burst frequency, and active electrodes showed significantly reduced neural activity in the ENOs caused by the acute fentanyl treatment, but no significant changes were detected in the CNOs under the same treatment condition (**Figure 6C**).

**Figure 6.**
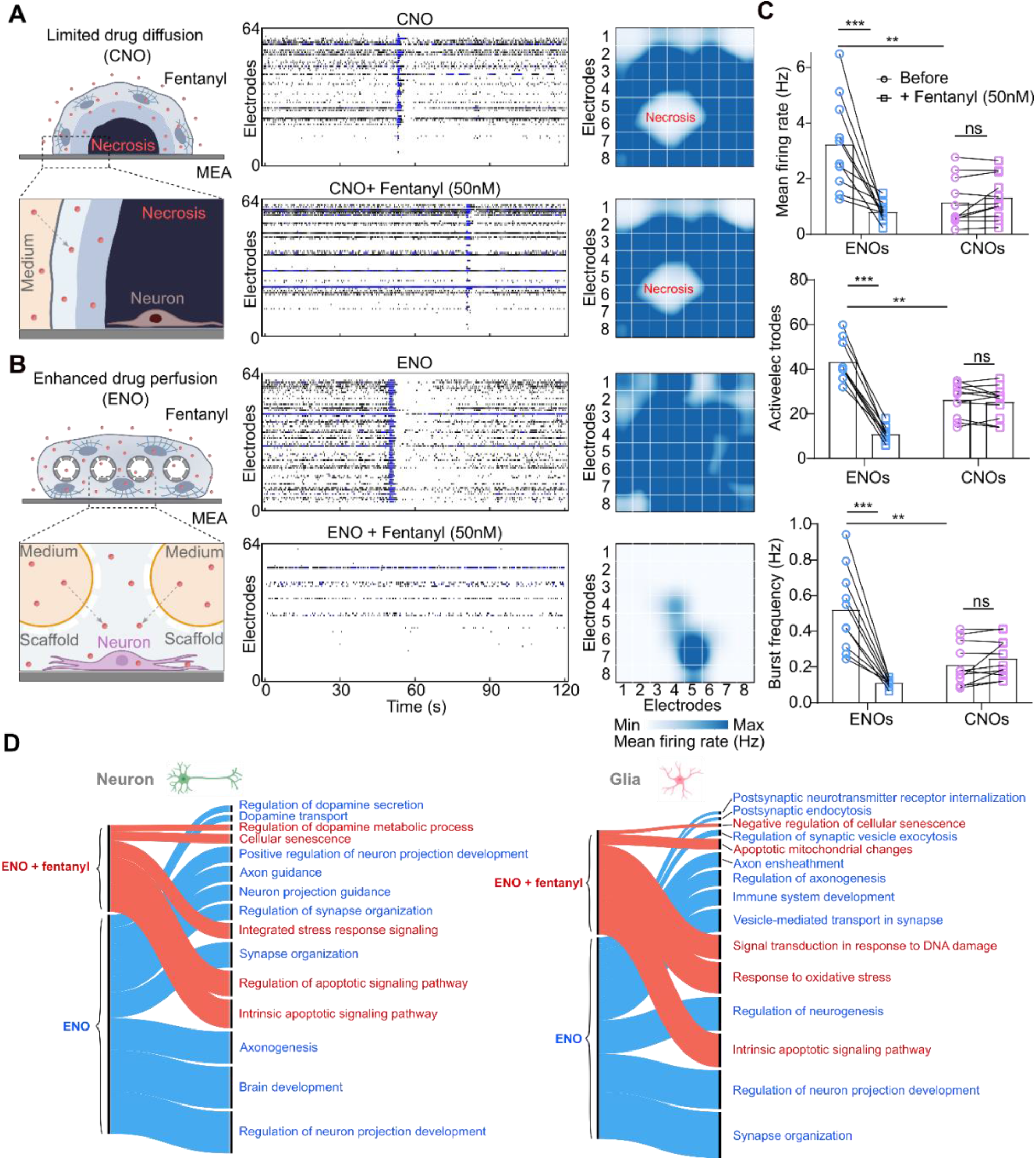
Enhanced pharmacological response. **(A)** Schematics of drug diffusion in the conventional neural organoid (CNO) on the microelectrode array (MEA) electrodes Representative raster plots and smoothed activity heatmaps of the same CNO before and 20 minutes after 50nM fentanyl treatment. **(B)** Schematics of drug diffusion in the engineered neural organoid (ENO) on the microelectrode array (MEA) electrodes. Representative raster plots and smoothed activity heatmaps of the same ENO before and 20 minutes after 50nM fentanyl treatment. **(C)** Quantification of electrical activity changes in response to fentanyl treatment, as indicated by mean firing rate, burst frequency, and active electrodes (mean± SEM, n = 11, from 3 independent experiments). **(D)** GO term analysis of the ENOs treated with 50nM fentanyl for 2 days (ENO + fentanyl) and the ENOs without any treatment (ENO). Alluvial plots show the gene ontology (GO) terms of neurons (Neuron) and astrocytes (Glia) in the ‘ENO’ and ‘ENO + fentanyl’ groups. The thickness indicates the number of significantly different gene expressions involved in each GO term.

To gain deeper insights into the effects of fentanyl treatment on the organoid neural networks, we conducted single-cell RNA sequencing analysis on the 2-month ENOs treated with 50nM fentanyl for 2 days (ENO + fentanyl) and the ENOs without any treatment (ENO). We focused on cell clusters of neurons (Neuron) and astrocytes (Glia), which are key components of organoid neural networks. Analysis of differentially expressed genes, followed by downstream Gene Ontology (GO) analysis (**Figure 6D**), revealed that genes upregulated in the neurons and astrocytes of the ‘ENO + fentanyl’ group were enriched in terms related to stress, apoptosis, and senescence, compared to the control group. In contrast, genes associated with dopamine processing, neuron development, and axon and synapse formation were significantly more highly expressed in the ‘ENO’ group compared to those in the ‘ENO + fentanyl’ group. Compared with CNOs, our ENOs demonstrated more physiologically relevant responses to fentanyl, presenting a promising platform for neural function-based compound and substance screening.

## DISCUSSION

We have designed and fabricated VID scaffolds that can fully recapture the benefits of physiological diffusion physics for organoid culturing when combined with orbital shaking-induced flows. We also demonstrated the successful generation and maturation of engineered neural organoids (ENOs), functional flattened human midbrain organoids with diffusible tube networks, from human stem cells. By supporting sufficient delivery of oxygen, nutrients, and other important growth factors during the whole developmental period, we showed that the ENOs have significantly improved cell survival, reduced cell stress, sustained neurogenesis, enhanced region-specific differentiation, and greater and more complex neural activity, compared to the conventional midbrain organoids. Finally, we also demonstrated an improved neural activity response to fentanyl in the ENOs, which was lacking in the CNOs. The scaffolds may offer a promising means for engineering and phenotyping functional organoids under physiological perfusion-like conditions.

Our VID scaffolds and engineered diffusible organoids have several advantageous features. First of all, the scaffolds can fully recapitulate the diffusion physics of physiological vascular networks for supplying sufficient oxygen, nutrients, and signaling molecules throughout the whole development and maturation process, avoiding inadequate profusion issues that detrimentally affect cell survival, activity, and function at all organoid developmental stages, a feature that has been difficult to achieve to date. These physiological perfusion-like conditions produce more physiology-relevant differentiation and functional maturation in the ENOs compared to CNOs, providing an important strategy for generating standardized organoids. Second, the scaffolds are biocompatible plastic devices fabricated using robust 3D printing, enabling unprecedented survival, development, and function of organoids over months. Third, the unique flattened shape and rigid material of the 3D printed scaffolds can guide organoid growth into a flattened shape keeping most of the organoid cells within diffusion limit (e.g., maximum D_nds_ < 150 μm) for nutrients. Meanwhile, the engineered flattened organoids with a thickness of less than 800 μm also support various characterization methods incorporating diverse applications because the flattened organoids are compatible with the flat MEA chip and the confocal imaging. Fourth, the dimensions of the scaffolds such as scaffold size, thickness, and tube network density can be adapted to fit with commonly used 24, 96, or 384 well-plates and are easily compatible with various organoid protocols in common lab settings. Fifth, the scaffolds can be massively produced using simple and inexpensive 3D printing, highlighting the potential for scalable, reproducible, and cost-effective generation of functional organoids for broad applications. Finally, the engineered diffusible functional organoids also reproduce a classical pharmacological response, highlighting their potential for drug screening. Together, these features make the 3D-printed scaffolds a promising means for engineering and characterizing various types of functional organoids.

## AUTHOR CONTRIBUTIONS

H.C. conceived and designed the study; performed the 3D printed scaffold fabrication, organoid culture, and characterization; conducted data analysis and interpretation; and wrote the manuscript. C.T. and L.C. performed the immunostaining and electrophysiology for organoid characterization. K. M., J. T., M. Gu., and K. M., performed data interpretation and edited the manuscript. F.G. conceived and designed the study, acquired the funding, supervised the study, performed data analysis and interpretation, and wrote the manuscript.

## ACKNOWLEDGMENTS

We thank Y. Miao for technical assistance. This study was supported in part by the National Institute of Health Awards (DP2AI160242 and U01DA056242). We also acknowledge the Indiana University Imaging Center (NIH1S10OD024988-01).

